# Oxytocin receptor activation does not mediate associative fear deficits in a Williams Syndrome model

**DOI:** 10.1101/2021.03.29.437431

**Authors:** Kayla R. Nygaard, Raylynn G. Swift, Rebecca M. Glick, Rachael E. Wagner, Susan E. Maloney, Georgianna G. Gould, Joseph D. Dougherty

**Author notes:** Corresponding Author: Joseph D. Dougherty, Depts of Genetics and Psychiatry, Washington University in St. Louis, 660 S. Euclid Ave, Campus Box 8232, St. Louis, MO 63110.

## Abstract

Williams Syndrome is caused by a deletion of 26-28 genes on chromosome 7q11.23. Patients with this disorder have distinct behavioral phenotypes including learning deficits, anxiety, increased phobias, and hypersociability. Some studies also suggest elevated blood oxytocin and altered oxytocin receptor expression, and this oxytocin dysregulation is hypothesized to be involved in the underlying mechanisms driving a subset of these phenotypes. A ‘Complete Deletion’ mouse, modeling the hemizygous critical region deletion in Williams Syndrome, recapitulates many of the phenotypes present in humans. These Complete Deletion mice also exhibited impaired fear responses in the conditioned fear task. Here, we address whether oxytocin dysregulation is responsible for this impaired associative fear memory response. We show direct delivery of an oxytocin receptor antagonist to the central nervous system did not rescue the attenuated contextual or cued fear memory responses in Complete Deletion mice. Thus, increased oxytocin signaling is not acutely responsible for this phenotype. We also evaluated oxytocin receptor and serotonin transporter availability in regions related to fear learning, memory, and sociability using autoradiography in wild type and Complete Deletion mice. While we identified trends in lowered oxytocin receptor expression in the lateral septal nucleus, and trends towards lowered serotonin transporter availability in the striatum and orbitofrontal cortex, we found no significant differences after correction. Together, these data suggest the fear conditioning anomalies in the Williams Syndrome mouse model are independent of any alterations in the oxytocinergic system caused by deletion of the Williams locus.

## INTRODUCTION

Williams Syndrome (WS), a multisystemic neurodevelopmental disorder, is caused by a 1.5—1.8 Mbp hemizygous deletion on chromosome 7q11.23, altering the copy number of 26-28 contiguous genes in the WS critical region (WSCR). The complex phenotypic characteristics of WS include craniofacial dysmorphology, connective tissue abnormalities, and cardiac problems such as supravalvular aortic stenosis and peripheral artery stenosis. In addition, WS is characterized by distinct cognitive features, including intellectual disability, profoundly impaired visuospatial construction,^1^ atypical facial processing, deficits in motor coordination and control, odynacusis,^2^ and impaired auditory processing.^3–12^ Interestingly, however, individuals with WS possess relatively intact expressive language and verbal skills,^13,14^ as well as heightened sensitivity and emotional response to music.^2,15^ One of the most striking phenotypes of individuals with WS is hypersociability and strong social motivation,^16–18^ despite high non-social anxiety^19^ and deficits in social cognition and awareness.^20^

A substantial body of research indicates the neuropeptide oxytocin (OT) plays a key role in mediating the regulation of social behavior and cognition, fear conditioning and extinction, observational fear,^21^ fear modulation via social memory,^22^ and anxiety in humans and rodents.^23–25^ Given the aberrant social behavior and anxiety in individuals with WS, recent studies have tested the hypothesis that OT is dysregulated in WS. Indeed, one study found elevated blood levels of OT in individuals with WS compared to controls.^26^ However, the findings on the oxytocin receptor (OXTR) have been contradictory. One study suggested increased gene expression,^27^ while another demonstrated downregulation and hypermethylation of *OXTR* in WS.^28^

While we did not see altered social behavior in a recent application of the standard social approach task, we did see differences in freezing during a conditioned fear task.^29^ Another mouse model, which deletes the entire WS-homologous region, has also shown alterations in fear conditioning,^30^ and individuals with WS have heightened phobias and non-social anxieties. Alterations in the brain OT system play an important role in social fear conditioning, contextual fear-induced freezing, and social fear extinction.^31,32^ Additionally, peripheral administration of an oxytocin receptor agonist has been shown to inhibit fear-induced freezing,^33^ and evoked oxytocin release via channelrhodopsins also results in attenuation of fear.^34^

In this study, we investigated whether OT dysregulation is a mechanism underlying the fear conditioning phenotype following deletion of the WSCR using the mouse experimental system. We employed the model reflecting the most common deletion found in WS patients: the hemizygous loss of the entire genomic region between the *Gtf2i* and *Fkbp6* genes.^30^ These heterozygous Complete Deletion (CD) mice show reduced freezing in fear conditioning recall, which is consistent with the expected consequences of OT elevation. Therefore, we probed whether OT activity could be responsible for the decreased expression of associative fear memory in CD mice. Further, to complement the prior human studies of OXTR expression in peripheral cells,^27,28^ we tested whether OXTR expression differs in CD versus wild type (WT) mice across the brain, but we found no differences after statistical correction in this system, nor in a second neurotransmitter system (serotonin, 5HT), which had previously been shown to cooperate with OT in social learning,^25^ and can be influenced by OT.^35^ Together, these data suggest there is not a direct role for the OT system in associative fear learning in WS.

## MATERIALS & METHODS

### Animals

CD mice contain a hemizygous deletion of the WSCR and were maintained on the C57BL/6J background (Jackson #000664).^30^ Animals were bred by crossing CD heterozygotes to C57BL/6J WT animals to produce heterozygous CD experimental mice along with WT littermates for the control group. Tissue collection and genotyping PCR occurred in the second postnatal week. Mice were housed by sex and treatment, when relevant, and were kept on a 12:12 h light/dark schedule with food and water provided ad libitum. All studies were approved by and conducted in accordance with the Institutional Animal Care and Use Committee at Washington University in St. Louis. All behavioral testing occurred during the light phase and was conducted by a female experimenter. Four independent cohorts were used in this study. Cohort 1 included 13 CD and 11 WT male mice from 8 independent litters, and was used to assess blood OT levels via ELISA. Cohort 2 comprised 10 CD (4 females (F), 6 males (M)) and 10 WT (8 F, 2 M) mice from four independent litters and were behaviorally examined as adults (postnatal day (P) 97—106) with the conditioned fear task. Cohort 3 comprised 14 CD (8 F, 6 M) and 29 WT (12 F, 17 M) mice from 16 independent litters, and served to evaluate the role of the OT system in associative fear and avoidance learning as adults (P68—118). Cohort 4, comprising 22 WT (12 F, 10 M) and 14 CD (5 F, 9 M) from 11 independent litters, was used to evaluate OXTR and serotonin transporter (SERT) expression in specific brain regions of interest. Tissue was collected post-mortem to confirm initial genotyping results.

### Oxytocin ELISA

Blood was drawn from the retro orbital sinus of isoflurane-anesthetized mice at P30 using heparinized glass capillary tubes. Samples were collected in 1.8 mL EDTA-coated tubes, spun at 1600 g for 5 min at 4 °C, then split into two aliquots, placed on dry ice, and stored at −80 °C until use. An ELISA kit was used for colorimetric quantification of OT per the manufacturer’s protocol (ADI-900-153A, Enzo Life Sciences, Farmingdale, NY). Prior to use, samples were diluted 1:3 with 200 μl of assay buffer. Absorbance measurements were read at 405 nm and OT concentration was calculated using a standard curve produced using the provided OT standards.

### Conditioned fear task

Associative fear and avoidance learning were evaluated in the CD mice using the conditioned fear paradigm (Figure 1A), as described in our previous studies.^36^ Briefly, each mouse was habituated to and tested in an acrylic apparatus, which measured 26 cm x 30 cm x 30.5 cm tall and contained a metal grid floor, an LED light bulb, and an inaccessible peppermint odorant, housed within a sound-attenuating chamber (Actimetrics, Wilmette, IL). The chamber light turned on at the start of each trial and remained illuminated for the duration. On Day 1, the testing session was five min. An 80 dB white noise tone sounded for 20 sec each at 100 sec, 160 sec and 220 sec. A 1.0 mA shock was paired with the last two sec of the tone. The baseline freezing behavior (first two min) and freezing behavior during the last three min was quantified via the computerized image analysis software program FreezeFrame (Actimetrics). This measure allowed for simultaneous visualization of behavior while adjusting a “freezing threshold,” which categorized behavior as freezing or not freezing during 0.75 sec intervals. Freezing was defined as no movement except for normal respiration, and data were presented as percent of time spent freezing. Testing on Day 2 lasted eight min, during which no tones or shocks were presented. This procedure enables evaluation of freezing behavior in response to contextual cues associated with the shock stimulus from Day 1. For the 10 min testing session on Day 3, the context was changed to an opaque Plexiglass-walled chamber containing a different (coconut) odorant. The 80 dB tone began at 120 sec and lasted the remainder of the trial. Freezing during habituation to the new context was quantified across the first two min. Freezing behavior to the auditory cue associated with the shock stimulus from Day 1 was quantified for the remaining eight min. Each day of testing, males were run first, followed by females. Assigned boxes were counterbalanced by genotype. Between animals, the apparatus was cleaned with 70% ethanol (Days 1 and 2) or 0.2% chlorhexidine diacetate solution (Day 3; Zoetis, Parsippany-Troy Hills, NJ). Animals were put in a holding cage until all cagemates had been tested, then animals were returned to their home cage. Shock sensitivity was evaluated after testing as previously described to verify differences in freezing were not the result of altered sensitivity to the shock stimulus itself.^37^

**FIGURE 1.**
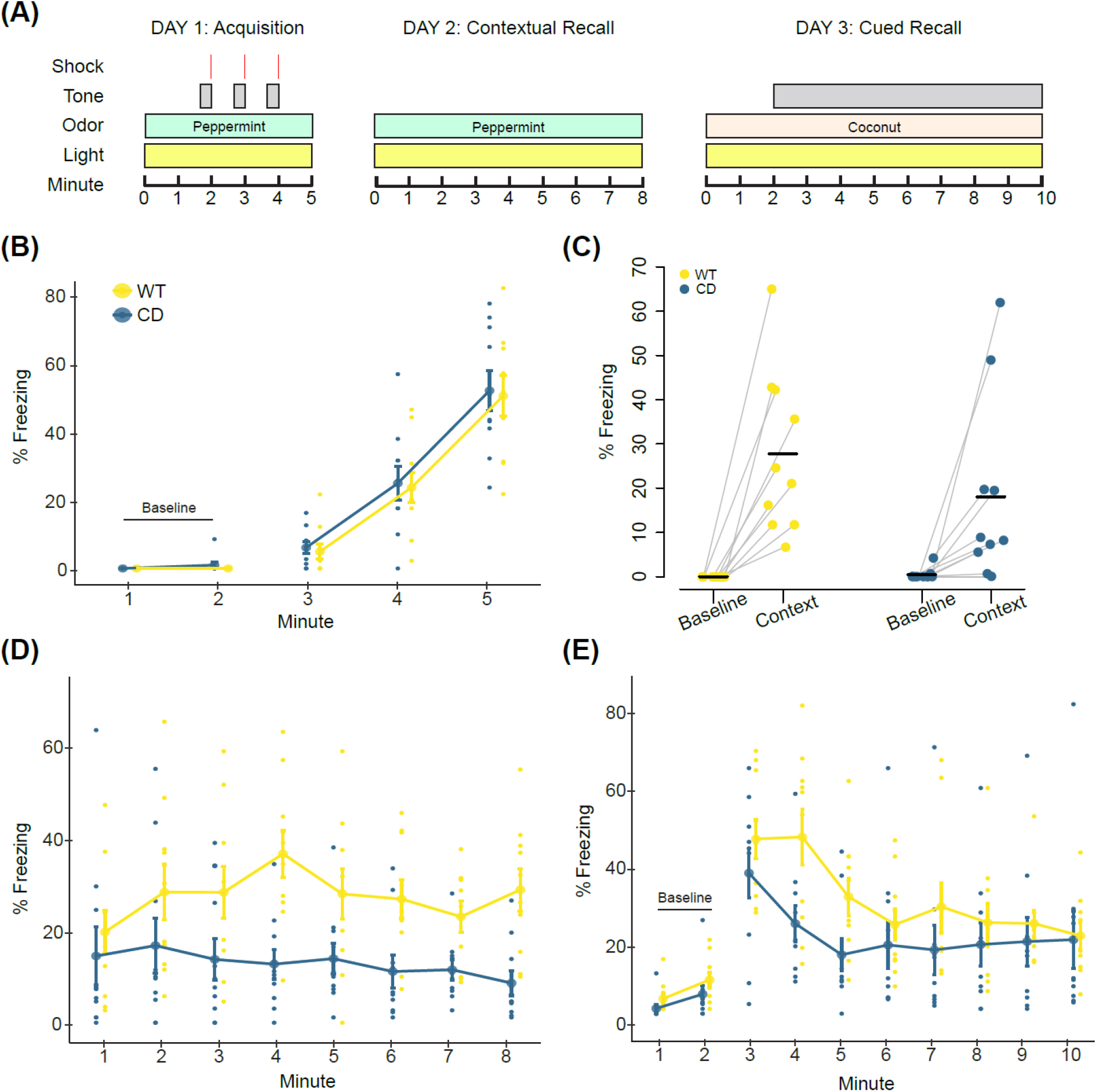
Complete Deletion mice have altered associated fear responses in a conditioned fear task. **A)** Overview of the conditioned fear task protocol. **B)** Day 1. CD and WT mice show increased freezing with subsequent footshock deliveries (main effect of minute, F(2,36)=7.8, p=8.5×10^-14^). **C)** All mice increased freezing in the context associated with the footshock (main effect of minute, F(1,18)=26.66, p=6.5×10^-5^). Baseline is the average of the first two minutes of Day 1. Context is the average of the first two minutes of Day 2. Black bars indicate the mean average percent freezing. Data points of individual mice are connected. **D)** Day 2. All mice increased freezing to context relative to Day 1 baseline, though CD mice freeze less than WT mice (F(1,18)=7.95, p=0.011). **E)** Day 3. CD mice have significantly decreased freezing relative to WT mice during minute 4 of tone delivery (p=0.036; trending minute by genotype interaction, F(9,162)=1.92, p=0.053). WT: n=10; CD: n=10. Connected data points in **B**, **D**, and **E** are means ± SEM. Individual scores are represented by colored circles in the background.

### Intracerebroventricular infusion of oxytocin receptor antagonist during conditioned fear task

The surgical area of adult mice was shaved a day prior to insertion of the guide cannula to facilitate the intracerebroventricular (ICV) injections. Mice were anesthetized with 2.5-5% isoflurane and placed in a stereotaxic apparatus. Prior to the procedure, mice received a local anesthetic, 1 mg/kg of Buprenorphine SR (ZooPharm, Laramie, WY), and an antibiotic, 2.5-5 mg/kg of Baytril (Bayer Healthcare LLC, Shawnee Mission, KS). An incision was made along the skull to visualize bregma to lamda. The periosteum was removed by lightly scratching the surface of the skull and the area was cleaned three times with a betadine solution (Purdue Products L.P., Stamford, CT) on sterile cotton swabs followed by a quick hydrogen peroxide swab. The guide cannula was placed in a stereotaxic cannula holder (Stoelting, #51636-1). Using a rapid, fluid motion, the 26-gauge unilateral guide cannula (C315GS-5/SPC, Plastics One, Roanoke, VA) with dummy cap was inserted at the following coordinates: M/L=+1, A/P=-0.4, D/V=-2.2, based on prior work.^38–40^ The guide cannula was cut to a length of 2mm so that it entered the lateral ventricle. The dummy cap (C315DCS-5/SPC) and internal cannula (C315IS-5/SPC) were cut to protrude 0.2mm from the end of the guide. C&B Metabond dental cement (Parkell, Edgewood, NY) was mixed on a chilled ceramic dish and used to secure the cannula to the skull and seal the surgical area. The dental cement dried completely before the animal was removed from the stereotaxic apparatus and placed in a recovery cage. Animals were housed together after fully awake and provided 0.25 mg of the chewable anti-inflammatory Rimadyl (Bio-Serv, Flemington, NJ). During daily monitoring, dummy caps were replaced and tightened as needed. Mice were euthanized at the first sign of distress or damage to the surgical area and had at least 3 days for recovery prior to testing.

All mice received 1 μl infusions at least one hour before each day of the conditioned fear task. Each mouse was given either vehicle (artificial cerebrospinal fluid solution, Tocris Bioscience, Bristol, UK; WT n=16, CD n=7) or an oxytocin receptor antagonist (OTA) (desGly-NH_2_,d(CH_2_)_5_[Tyr(Me)^2^Thr^4^]OVT, Bachem, Torrance, CA; WT n=13, CD n=7). The OTA, dissolved in vehicle at 1 ng/uL, is a peptidergic ornithine vasotocin analog chosen because of its broad applicability and prior use in ICV injections.^41,42^ The solutions were unilaterally delivered into the lateral ventricles through the 33-gage internal cannula via a PlasticsOne Cannula Connector (C313CS) over the course of one min using a Quintessential Stereotaxic Injector (Stoelting #53311) and a 1 μl Hamilton syringe. After injection, 15-30 sec passed before removing the internal cannula to ensure proper diffusion. The conditioned fear task was performed as described above. Following completion of behavioral testing, cannula placement was confirmed by injecting enough dye to flood the ventricles and immediately euthanizing the animal via isoflurane overdose. Brains were extracted and sliced coronally at the injection site with a razor blade. Infusion of the dye into the ventricles was then confirmed by eye and samples that missed the ventricles were excluded from the final analysis.

### Quantitative autoradiography of OXTR and SERT in mouse brain

Naive mice were rapidly euthanized by cervical dislocation. Brains were removed, placed in an ice cold saline solution for one min, then excess saline solution was wicked onto a paper towel. Brains were frozen on crushed dry ice and then stored at −80 °C until sliced into 20 μm coronal sections in a cryostat (Leica Biosystems 1850, Buffalo Grove, IL) at −16 to −18 °C. Slides were thaw-mounted onto gelatin-coated microscope slides, vacuum-desiccated overnight (18 h) at 4 °C, then stored at −80 °C until use. Adjacent sections were used for two distinct ligands, [^125^I]OVT and [^125^I]RTI-55, to assess OXTR and SERT availability, respectively.

#### OXTR Quantitative Autoradiography

Binding of iodinated ornithine vasotocin analog ([^125^I]OVT) to OXTR in the mouse brain was performed as described previously,^43^ with minor modifications. Mounted sections were thawed for 30 min at 22-23 °C, then pre-incubated for 30 min in 50 mM Tris HCl buffer pH 7.4 at 22-23 °C. Next, sections on slides were incubated for 90 min in upright cytomailers filled with 10 ml buffer containing 10 mM MgCl_2_, 0.1% bovine serum albumin and 50 pM [^125^I]OVT (NEX2540, PerkinElmer, Boston, MA). Non-specific binding was obtained by incubating representative adjacent sections on slides from the series in buffer containing unlabeled OT (1 μM, Ascent Scientific, Bristol, UK). Sections on slides were then washed twice for five min each in glass staining dishes containing 300 ml of 4 °C buffer, and were dipped for two sec in 4 °C deionized water. Slides were dried on a benchtop slide warmer for one hour or until sections were opaque and any droplets had evaporated.

#### SERT Quantitative Autoradiography

Slides with brain sections were defrosted at 22-23 °C for 30 min and pre-incubated in 30 mM sodium phosphate, 120 mM sodium chloride buffer, pH 7.4 at 22-23 °C, for 30 min. For incubation, 100 mM sucrose, 100 nM GBR12909 (to block binding to dopamine transporter) and 50 pM [^125^I]RTI-55 (NEX272, Perkin Elmer, Boston, MA) were added to the buffer. To measure non-specific binding, 10 μM mazindol was added to a subset of slide mailers containing a representative set of duplicate slides. All unlabeled ligands were from Sigma. Incubation was carried out for two hours at 22-23 °C. Sections were rinsed twice for one min in 4 °C buffer (without sucrose), then dipped for two seconds in 4 °C deionized water, drained and placed on a slide warmer (Lab-Line, Fisher Scientific, USA), at moderate setting (4 on 10 scale) for two hours.

#### Exposure and Imaging

Sections on slides were exposed to Biomax MR film (Carestream/Kodak) in a cassette for 48 h along with tritium standards (ART0123A, American Radiolabeled Chemicals, St. Louis, MO) calibrated to [^125^I]-incubated brain mash, as previously described.^44^ Films were developed using an automatic film processor. Digital images of autoradiograms were captured using a 12-bit CCD monochrome digital camera (CFW-1612M, Scion, Frederick, MD) with a 60 mm lens (f-stop = 4)(Nikon, Melville, NY) mounted on a copy stand (RS-1, Kaiser Fototechnik, White Plains, NY) with an LED lightbox (Slimlite Plano, Kaiser). Pixel intensity was calibrated to measure density in units of femtomoles/mg (fmol/mg) protein using a linear function with ImageJ software (https://imagej.nih.gov/ij/download.html).^45^

#### OXTR Data Collection

OVT binding to the OXTR was measured in the anterior olfactory nucleus (AON), lateral septal nucleus (LSN), anterior cingulate cortex (ACC), striatum (CPu), hippocampal CA2 and CA3 regions (CA 2/3), paraventricular hypothalamic nucleus (PVN), piriform cortex (Pir), and the combined basolateral and lateral amygdala (BLA & LA) by tracing each region in ImageJ based on coordinates from the Franklin and Paxinos mouse brain atlas.^46^ The AON was traced at Bregma 2.68 mm. The LSN, ACC, and CPu were traced between Bregma 0.68 mm and 0.26 mm. The hippocampal CA 2/3 region, PVN and BLA & LA were traced between Bregma −1.58 mm and −1.82 mm. Each region was measured at least twice in nine or more animals per genotype (Supplemental Table 2).

#### SERT Data Collection

SERT availability was measured in the BLA, LA, central amygdala (CeA), ACC, CPu, CA 2/3, insular cortex (IC), lateral parietal association (PtA), nucleus accumbens (NAc), orbitofrontal cortex (OFC), peduncular part of the lateral hypothalamus (PLH), and the bed nucleus of the stria terminalis (BNST). The BLA, LA, and CeA were traced between Bregma −1.58 and −1.82 mm. The ACC was traced at Bregma 0.74 mm, the IC was traced at Bregma −1.06 mm, and the NAc was traced at 0.74 mm. The CPu was traced between Bregma 0.74 mm and 0.26 mm. The hippocampal CA 2/3 region was traced between Bregma −1.34 mm and −1.58 mm. The PtA and PLH were traced at Bregma −1.58 mm to −1.70 mm. The BNST was traced between Bregma 0.62 mm and −0.22 mm. Up to four independent measurements were taken within a sample and at least five animals were measured per genotype (Supplemental Table 2).

### Statistical analysis

Analysis for ELISA and the conditioned fear task was completed in R using RStudio (Version 1.2.5019). ELISA analysis utilized the ‘drc’ package.^47^ Conditioned fear data were condensed by minute then assessed for normality, homogeneity of variance, and outliers. Data were analyzed with a linear mixed effect model using the ‘lme4’ package,^48^ with genotype as the main factor, and minute as a repeated measure. Tukey’s HSD was employed for post hoc analysis. An effect of sex was screened for but not included in the final analysis, as there was no significant effect on any outcome. Detailed outputs are included in Supplemental Table 1.

Autoradiography analysis was performed using IBM SPSS Statistics for Windows, Version 26.0. Measurements within a region were averaged to compute the binding value of that region for each sample. These values were normalized by film to compute the total binding value for each region by subtracting the nonspecific binding value of a control sample from the sample binding value. Values less than zero after normalization were treated as zero. Normality, outliers, and variance were assessed prior to hypothesis testing. Total OXTR binding values were transformed using a square root transformation to resolve normality violations. A 2×2 Analysis of Covariance (ANCOVA) with fixed factors of sex and genotype and a covariate of age was performed on each region of interest for OXTR autoradiography. For SERT, no transformations were necessary. A 2×2 multivariate ANOVA was used assessing the factors of sex and genotype across 12 regions of interest. Detailed outputs of statistical tests are in Supplemental Table 2.

## RESULTS

### Complete deletion mice show impaired contextual and cued fear conditioning

OT has been shown to modulate the freezing response of rodents in conditioned fear tasks. Specifically, central administration of OT results in decreased levels of freezing in response to avoidance-associated cue or context.^49^ Thus, due to the suggested increase in OT production in WS, we sought to determine whether the CD mouse model also had altered associative fear learning. Previously a decrease in overall freezing time during fear conditioning was shown in the CD model using only male mice;^30^ here we replicate and expand on these findings by confirming the phenotype in both sexes.

We found that CD mice responded to shock during conditioning (Day 1) by increasing freezing to the same extent as WT mice with a main effect of minute (F(2,36)=77.81, p=8.5×10^-14^), but no main effect of genotype or minute by genotype interaction (Figure 1B, complete statistical analysis available in Supplemental Table 1). WT and CD mice both exhibit significantly increased freezing (F(3,36)=9.6, p=8.5×10^-5^) in the first two minutes of the contextual memory test (context) compared to the first two minutes of training (baseline), indicating each group successfully associated the fear stimuli with the context (Figure 1C). On Day 2, CD mice froze less compared to WT mice when placed in the same context (chamber/odor) used for fear conditioning (Figure 1D, main effect of genotype, F(1,18)=7.95, p=0.011), indicating impaired contextual associative fear memory. The response to the conditioned cue was also decreased in CD mice but not as broadly, with a main effect of minute (F(9,162)=22.0195, p=2×10^-16^), and a borderline minute by genotype interaction (Figure 1E, F(9,162)=1.9151, p=0.053), but no main effect of genotype. Post hoc analysis revealed the mean freezing of CD mice was almost 24% less than WT freezing at minute 4 (p=0.036). Overall, CD mice show a deficit in contextual fear conditioning that is consistent with elevated OT activity.

### CD conditioning deficits are not reversed by central infusion of an oxytocin receptor antagonist

We next sought to determine if CD mice had elevated OT levels in the blood, as had been reported in patients.^26^ We used the same ELISA approach, but did not see a significant difference between genotypes (t=0.003, df=21.664, p=0.9976; CD (n=13): *M*=1138.191 pg/mL, *SD*=503.682; WT (n=11) *M*=1138.791 pg/mL, *SD*=478.916). However, even WT mice had a remarkable range of OT levels in blood (82-1731 pg/mL), likely driven by the periodic nature of OT release, coupled with its short half life (<5 min).^50^ Thus, it can be hard, at a single point in time, to detect and make conclusions about average OT levels. Therefore, we took an experimental approach; if elevated OT was responsible for decreased associative fear learning in CD mice, then blocking OXTR activity should reverse the phenotype. We implanted CD and WT mice with ICV cannulas to directly administer an OXTR peptide antagonist (OTA) and block all receptor activity during the fear conditioning procedure (Figure 2A).

**FIGURE 2.**
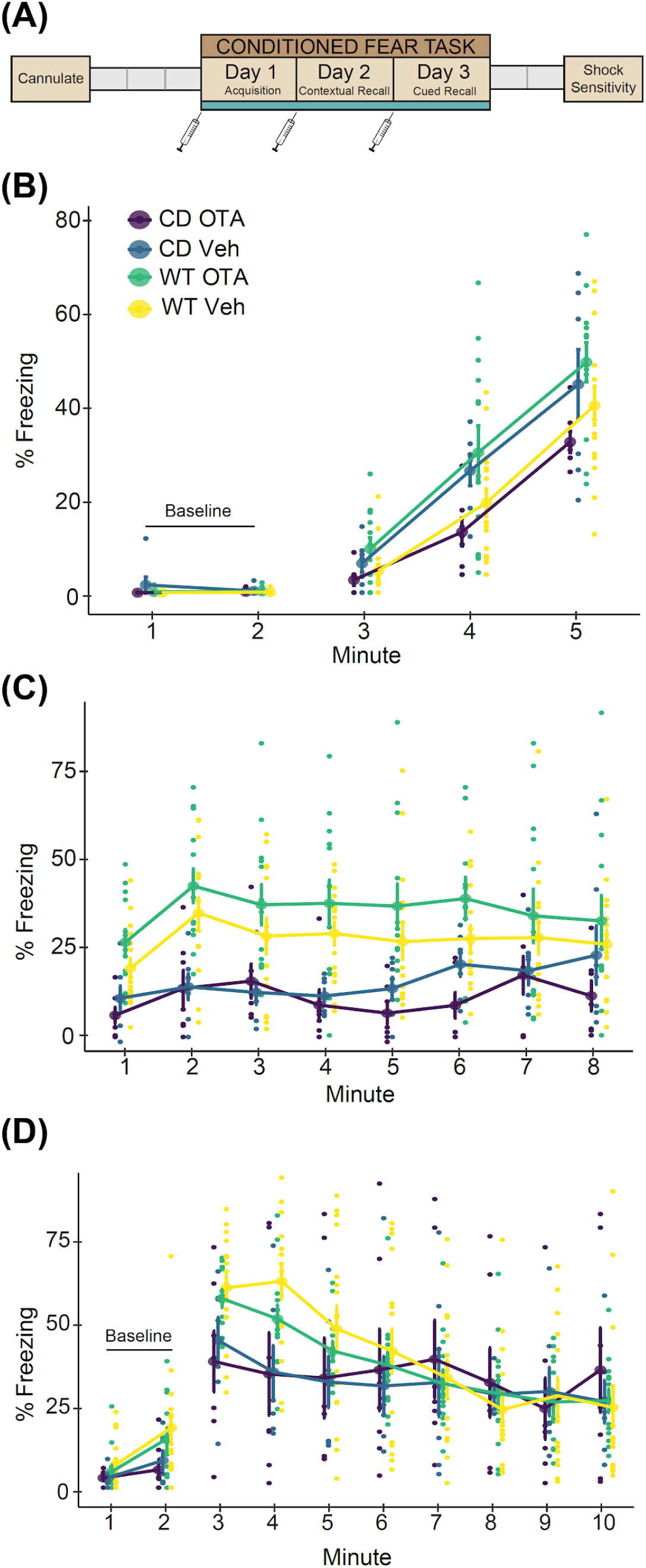
Central administration of an oxytocin receptor antagonist does not rescue reduced contextual or cued fear responses in CD mice. **A)** Schematic of the experiment. Grey boxes show rest days. Syringes indicate ICV infusions of OTA or Vehicle, which occurred at least 1 hour prior to testing. **B)** Day 1 of conditioned fear. CD and WT mice show increased freezing with subsequent footshock deliveries (main effect of minute, F(2,78)=155.18, p=2×10^-16^). WT OTA-treated mice freeze significantly more than CD OTA-treated mice (interaction of genotype and treatment, F(1,39)=7.32, p=0.01). **C)** Day 2. All mice show increased freezing to context (minutes 1 and 2) relative to Day 1 baseline (main effect of minute, F(1,39)=196.3, p=2.2×10^-16^). CD mice freeze less than WT mice (main effect of genotype, F(1,39)=19.96, p=6.6×10^-5^) and there is no main or interaction effect of treatment. **D)** Day 3. CD mice have significantly decreased freezing relative to WT during minute 4 of tone delivery (p=0.008; minute by genotype interaction, F(7,273)=5.87, p=2.26×10^-6^), but there is no effect of treatment. WT Veh: n=16; WT OTA: n=13; CD Veh: n=7; CD OTA: n=7. Connected data points are means ± SEM. Individual scores are represented by smaller unconnected circles. Veh = vehicle-treated; OTA = oxytocin receptor antagonist-treated.

There were no baseline freezing differences between CD and WT animals, as measured in minutes 1 and 2 of the first day of testing (F(1,39)=1.58, p=0.22). During the conditioning phase, we found an interaction of genotype and treatment (Figure 2B, F(1,39)=7.32, p=0.0101). This interaction results from the OTA having an opposite effect on freezing in CD and WT animals. Specifically, WT mice receiving the OTA froze more than their CD counterparts, with trending significance in post hoc analysis at minute 4 (p=0.078) and minute 5 (p=0.077).

During contextual fear recall, both CD and WT animals showed evidence of learning, as freezing increased in all groups from Day 1 baseline compared to the first two minutes of Day 2. Overall, a main effect of genotype on contextual fear memory (Figure 2C, F(1,39)=19.96, p<6.6×10^−5^) reflects a significant reduction in freezing within CD mice compared to WT mice, regardless of treatment. While this replicated results from our first experiment, there was no main effect of treatment or a genotype by treatment interaction, thus administration of the OTA did not significantly alter contextual fear responses. This was also true on Day 3 for cued fear responses (Figure 2D), where there was only a main of effect of time (F(2,273)=26.45, p=2.2×10^-16^) and an interaction between time and genotype (F(2,273)=5.88, p=2.26×10^-6^) driven by the decreased freezing of CD mice compared to WT mice at minute 4 (p=0.0083). Thus, we show OT signaling does not account for the impaired associative fear response in this model.

### Autoradiography reveals no changes in oxytocin receptor density or distribution in CD mice

In parallel to the experimental approach, we investigated OXTR availability in the mouse brain through a discovery-based approach, as elevated levels in the amygdala might influence fear conditioning.^49^ Particularly, given opposite directions of effect of the OTA in WT and CD mice on Day 1 freezing (Figure 2B), we suspected genotype differences in OXTR expression in regions related to fear learning. Furthermore, it is of interest to study this binding given the findings of OXTR dysregulation in brains of humans with the WSCR deletion.^27,28^ Therefore, we conducted an autoradiography study using radiolabeled OVT ligand on coronal sections of WT and CD brains (Figure 3A). As well as measuring regions relevant to fear conditioning (BLA and LA;^51,52^ LSN^31^), we measured areas of where OT has been shown to affect sociability or memory (AON;^53^ LSN;^54^ ACC;^55,56^ CA2/3;^57,58^ Pir;^59^ PVN^60^; and CPu^25^)(Figure 3B), as hypersociability and cognitive impairments are characteristic of WS. We found no significant differences in OXTR binding between genotypes within regions of interest in CD and WT brains when corrected for multiple testing (Figure 3C, Supplemental Table 2). There was a nominally significant change in the LSN (p=0.034), but it did not meet the corrected experimentwise critical alpha level (α=0.006). It is an interesting trend to report nonetheless, given the role of the LSN in fear and anxiety, and may motivate focused studies in the region.

**FIGURE 3.**
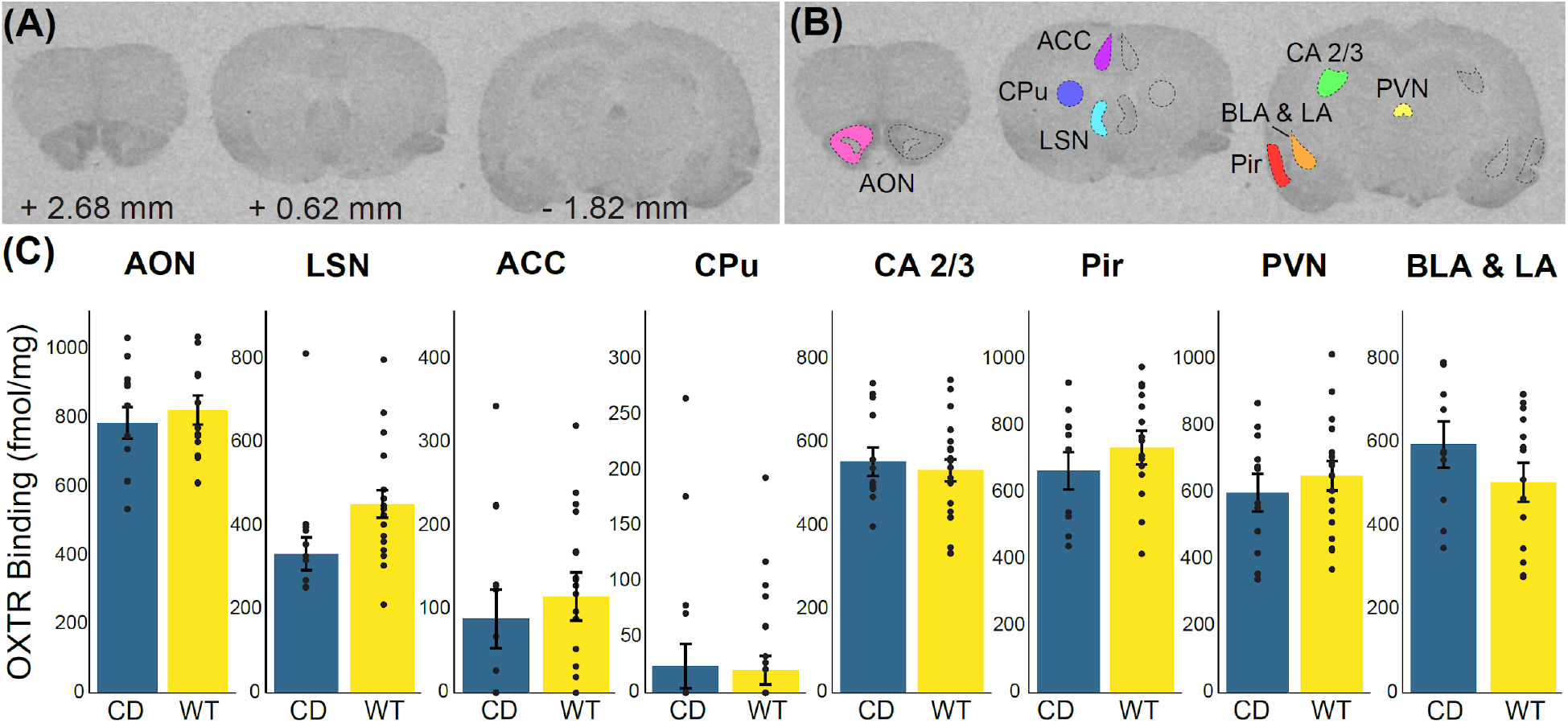
No significant differences in oxytocin receptor density in Complete Deletion mice. A) Coronal sections from a representative mouse brain used in iodinated ornithine vasotocin analog ([^125^I]OVT) autoradiography analysis with corresponding distance from bregma. **B)** Example tracing of regions of interest including anterior olfactory nucleus (AON), cingulate cortical areas 1 and 2 (ACC), striatum (CPu), lateral septal nucleus (LSN), hippocampal CA 2 and 3 regions, paraventricular nucleus (PVN), basolateral (BLA) and lateral (LA) amygdala, and piriform cortex (Pir). **C)** Results of oxytocin receptor autoradiography comparing [^125^I]OVT binding (fmol/mg of protein) in regions of interest between CD mice and WT mice. Colored bars show means ± SEM (brackets). No regions were significantly different after correction for multiple testing. Individual averaged measurements for each mouse are represented by circles. For each genotype, n ≥ 9.

**FIGURE 4.**
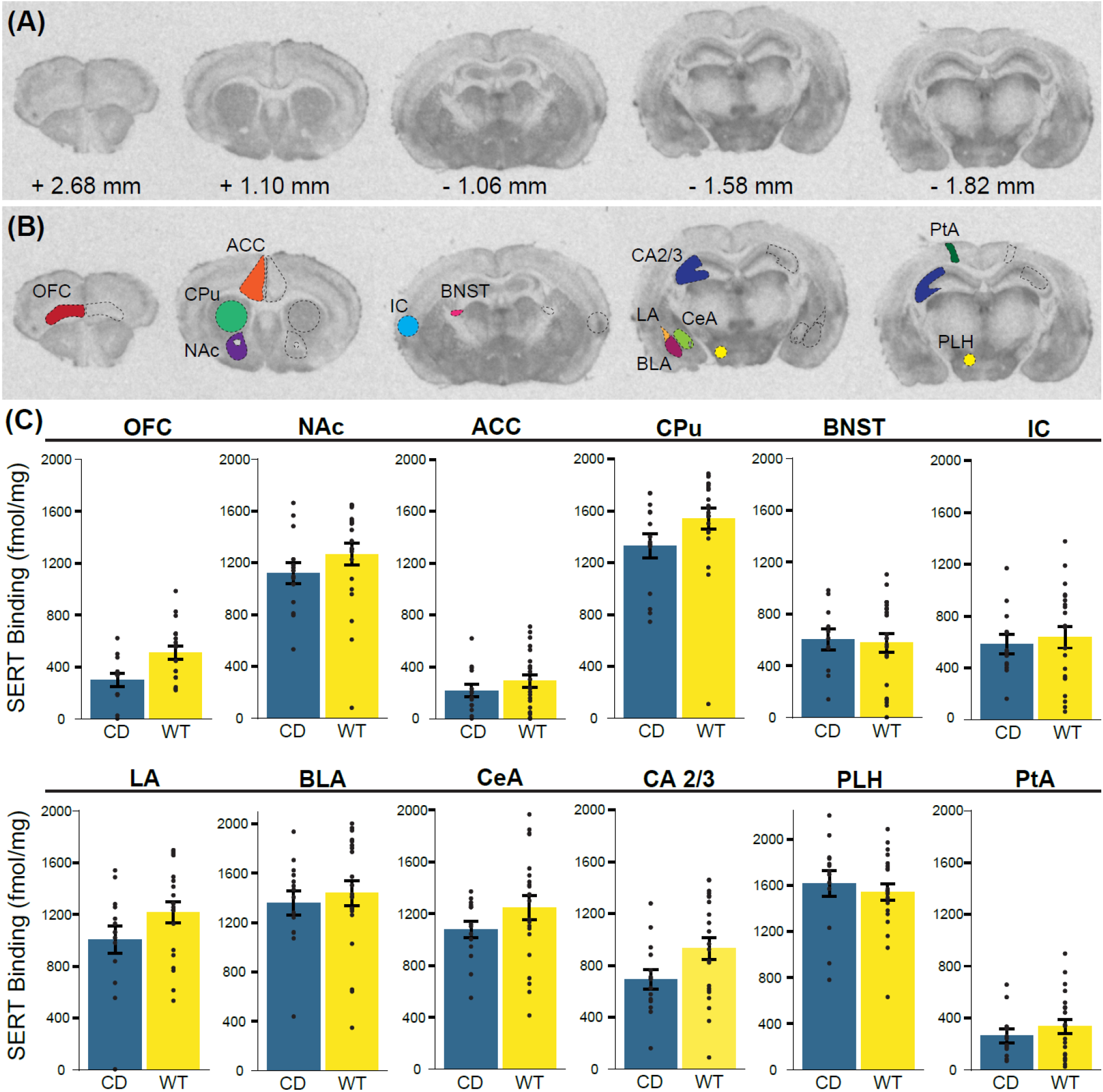
No significant differences in serotonin transporter density in Complete Deletion mice. A) Coronal sections with corresponding distance from bregma from a representative mouse brain used in [^125^I]RTI-55 autoradiography analysis. **B)** Example tracing of regions of interest including orbitofrontal cortex (OFC), anterior cingulate cortex (ACC), striatum (CPu), nucleus accumbens (NAc), the bed nucleus of the stria terminalis (BNST), insular cortex (IC), hippocampal CA 2 and 3 (CA 2/3) regions, basolateral (BLA), lateral (LA), and central (CeA) amygdala, lateral parietal association (PtA), and the peduncular part of the lateral hypothalamus (PLH). **C)** Results of serotonin transporter (SERT) autoradiography comparing [^125^I]RTI-55 binding (fmol/mg protein) in regions of interest between CD mice and WT mice. No regions were significantly different after multiple testing corrections, though multivariate ANOVA showed a trend towards a main of effect of genotype (F(1,22)=2.4; p=0.07); two individual regions were trending towards significance (CPu, p=0.067 and OFC, p=0.06). Colored bars show means ± SEM (brackets). Individual measurements for each mouse (regional average) are represented by circles. For each genotype, n ≥ 5.

### Autoradiography reveals no changes in serotonin transporter density or distribution in CD mice

Finally, with no changes in OXTR availability, we examined an additional alternative neurotransmitter: serotonin (5HT). Disruption to the 5HT system in WS has been suggested in prior studies. Specifically, Proulx et al. (2010) determined that there are enhanced 5-HT_1A_ receptor-mediated currents in a WS mouse model with low innate anxiety.^61^ More recently, Lew et al. (2020) compared serotonergic innervation in the amygdala between autism and WS in human postmortem samples, concluding that there is decreased innervation in WS brains compared to neurotypical brains.^62^ Therefore, we focused on the SERT, as it should provide a measure of serotonergic innervation to different structures.

We measured SERT binding in several brain regions. We focused on amygdalar regions relevant to the human postmortem studies^62^ and fear conditioning phenotypes,^63^ assessing BLA, central amygdala (CeA), and LA regions independently based on findings that they differ in the amount of serotonergic innervation in some species.^64^ We then included other areas where SERT has been implicated in behaviors relevant to the WS phenotype, patient findings, and knowledge of 5HT biology. These included the nucleus accumbens (NAc) based on 5HT’s role on social reward in this region,^25^ the BNST for its role in adaptive anxiety,^65^ and additional hypothalamic and cortical regions of interest, including the ACC, lateral parietal association (PtA), orbitofrontal cortex (OFC), and the peduncular part of the lateral hypothalamus (PLH) which is where the medial forebrain bundle is found, representing a major ascending pathway for nearly all 5HT axons.^66^ Overall, there was a trend towards a cross-region effect of genotype (F(12,12)=2.41; p=0.07), with no effect of sex (F(12,12)=0.49; p=0.89). In examining individual regions that might be driving this trend, we did not identify any regions with significant differences that survived multiple testing corrections, but trends of interest in the CPu (p=0.067) and OFC (p=0.060), suggest possible directions for future research.

## DISCUSSION

We and others have previously observed altered fear conditioning in WS models.^29,30^ Expanding on previous studies that used only male mice, we found, using both sexes, that CD animals on a C57BL/6J background had a suppressed fear response to the context and cue presented in the fear conditioning paradigm. Thus, CD mice enable investigation of the underlying circuit disruptions mediating this phenotype. We hypothesized that the altered associative fear memory response in CD mice was due to the increased availability of oxytocin based on human findings of elevated OT in WS.^26^ We focused our efforts on functional studies, which would be definitive with regards to a role for OT in fear conditioning of CD mice. Using an intraventricular cannula, we treated mice with an oxytocin receptor antagonist during each day of conditioned fear to attempt to counteract any possible effect of increased oxytocin production on the fear response. The OTA did not alter the subdued freezing response in CD mice. Therefore, the associative fear conditioning phenotype that results from loss of the WS critical region is not mediated by OXTR activity.

Our results tell us less about the role of OT in WT mice. While many prior studies show oxytocin modulating fear conditioning,^33,34,49,52,67^ Pisansky et al. (2017) found oxytocin enhances fear in a social paradigm of observational fear learning, but does not affect non-social fear learning.^21^ We did not see a significant difference (p<0.083, Day 2) in WT mouse behavior in response to OTA, but, as there was a trend, we are hesitant to weigh in on this debate. In the end, the discrepancy across studies could be a result of dosage. Gunduz-Cinar et al. (2020) show different concentrations of OT have opposing effects on other fear-related tasks.^68^ We have also observed an effect of genetic background on fear conditioning in the CD mice, with elevated freezing in CD mutants on an FVB/AntJ x C57BL/6J F1 hybrid background.^29^ These hybrids show different levels of baseline freezing in response to cues even in WT animals, suggesting the impact of the CD deletion interacts with other genes in the genome to modulate conditioned fear effects. Thus, if OT is not involved, a genetic screen for interacting loci using mouse strain panels may help identify the relevant pathways. Another option would be to investigate genes based on dysregulated expression in models of WS, such as the serotonin 5HT_1B_ receptor, which is among the top 10 dysregulated genes in a cell model of WS.^69^

In addition, given some of the prior work suggesting OXTR receptor gene expression and methylation in WS patient blood cells, we were curious if receptor availability was also altered in the brain following deletion of these genes. We used autoradiography because it can measure the availability of the receptor at the surface, which should reflect protein level and localization changes in addition to changes in gene expression. Further, it provides an opportunity for spatially informed analyses. As such, it is the best single measure for assessing if the OXTR is modulated in a conserved way by these mutations. We assessed regions previously associated with fear conditioning and those where OT had been shown to modulate social reward (CPu). Overall, we found no significant changes in OXTR binding in the CD mouse brain compared to controls, though there was a nominally significant difference in the LSN prior to correction, in the same direction of effect as seen in WS patient blood cells. The LSN is an interesting center integrating a variety of fear and anxiety signals, for example, playing a role in how stressful social cues are received.^70^ In addition, the LSN has been associated with fear-enhancing effects of the OXTR, but as modulation of the OXTR did not alter contextual conditioned fear responses, it is thought to be through indirect means.^71^ These data motivate future studies focusing on the role of the LSN in behavioral abnormalities observed in the CD mouse model.

Despite ruling out a direct role for OT in fear conditioning deficits in this model, altered OT might play a role in the increased social motivation in this population. While it is of interest scientifically to assess this, hypersociality is not as much of a concern therapeutically as other phenotypes, such as learning deficits, ADHD, phobias, and anxiety. Beyond oxytocin and serotonin, dopamine has also been implicated in WS,^7^ and has been previously connected to fear conditioning,^72^ other anxiety-avoidance tasks,^73^ and ADHD-related hyperactivity.^74^ Thus, it is possible deletion of the WSCR disrupts dopamine signaling to result in these behavioral alterations. Understanding the roles of any of these systems in patient-related phenotypes of the CD mice might help highlight potential treatments in WS, as a wide range of therapeutics working on these systems are currently available.

## ACKNOWLEDGEMENTS

We would like to thank N. Kopp for his contributions and support, E. Minakova and D. Burek for surgical training, and H.A. Stephens, Y.R. Sanchez, M.C. Nunn, and B.L. Pierce for their expert technical assistance in tissue preparation. Complete Deletion mice were a generous gift from V. Campuzano. The Animal Behavior Core at Washington University in St. Louis graciously shared their resources. Financial support for this work was provided by the NIMH 5R01MH107515-05 (JDD), NSF DGE-1745038 (KRN), NICHD P50HD103525 (IDDRC@WUSTL) and John L. Santikos Charitable Foundation of the San Antonio Area Foundation and The Morrison Trust (GGG).

